# CDC50A dependent phosphatidylserine exposure induces inhibitory post-synapse elimination by microglia

**DOI:** 10.1101/2020.04.25.060616

**Authors:** Jungjoo Park, Eunji Jung, Seung-Hee Lee, Won-Suk Chung

**Affiliations:** Department of Biological Sciences, Korea Advanced Institute of Science and Technology, Daejeon 34141, Republic of Korea

## Abstract

Glia contribute to synapse elimination through phagocytosis in the central nervous system. Despite important roles during development and neurological disorders, the “eat-me” signal that initiates glia-mediated phagocytosis of synapses remains largely elusive. Here, by generating inducible conditional knockout mice of *Cdc50a*, we induced stable exposure of phosphatidylserine in the neuronal outer membrane. Surprisingly, acute *Cdc50a* deletion in neurons causes specific loss of inhibitory post-synapses without affecting other synapses, thereby generating excessive excitability with appearance of seizure. Ablating microglia or deleting microglial *Mertk* rescues the loss of inhibitory post-synapses, indicating that microglial phagocytosis is responsible for inhibitory post-synapse elimination. Moreover, inhibitory post-synapses in normal juvenile brains also use phosphatidylserine for synapse pruning by microglia, suggesting that phosphatidylserine may serve as a general “eat-me” signal for inhibitory post-synapse elimination.

**One Sentence Summary:** *Cdc50a* dependent phosphatidylserine exposure functions as an “eat-me” signal for microglia-dependent inhibitory post-synapse elimination

## Main Text

Synapse elimination, the process of selectively removing unnecessary synapses, occurs in the central nervous system during development and adult stages as well as in various neurological disorders (*1*). Recent studies have shown that microglia and astrocytes contribute to synapse elimination through complement and MERTK/MEGF10 phagocytic pathways, respectively (*2-4*). Interestingly, complement-dependent microglial phagocytosis seems to participate in eliminating either excitatory or inhibitory synapses during development and diseased conditions, often creating excitatory-inhibitory imbalance (*3-7*). However, the identity of the “eat-me” signal that allows these glial phagocytic receptors to recognize specific synapses for subsequent elimination is still unclear. Moreover, whether excitatory and inhibitory synapses utilize the same or distinct mechanisms for presenting “eat-me” signals are completely unknown.

Phosphatidylserine (PS) is a phospholipid that predominantly resides on the inner plasma membrane under normal physiological conditions. When cells receive stress or apoptotic stimuli, PS flips out to the outer plasma membrane, functioning as an “eat-me” signal for subsequent engulfment by neighboring phagocytes (*8*). Among the upstream enzymes that control PS localization between the inner and the outer plasma membranes, type IV P-type Adenosine triphosphatases (P4-ATPases) have been recently identified as the flippases that use Adenosine Triphosphate (ATP) to translocate PS from the outer to the inner plasma membrane. Although many functionally redundant P4-ATPases are expressed in various tissues, most P4-ATPase family members require the activity of the flippase chaperone Cell Cycle Control protein 50A (CDC50A) for proper localization on the plasma membrane (*8, 9*). Previous *in vitro* studies have shown that a *Cdc50a* null mutation causes constitutive PS exposure in the outer plasma membrane without general apoptosis, and that PS exposure is sufficient to initiate phagocytic engulfment by macrophages (*9*). Although CDC50A is also highly expressed in mammalian brains (*10*), their potential roles in synapse elimination have not yet been studied.

To understand the function of CDC50A in synapse elimination, we first investigated whether *Cdc50a* is expressed in neurons from juvenile mouse brains. By using *Cdc50a* knockout-first allele heterozygous mice, which express a knocked-in *lacZ* gene at the *Cdc50a* locus, we found that β-Galactosidase was highly localized in both CAMKII-positive and GAD67-positive neurons (Fig. 1, A and B). When we injected adeno-associated virus (AAV) 9-*Cdc50a-mCherry* in wild-type (WT) mice brains, we found that CDC50A-mCherry was mostly localized in the neuronal rough endoplasmic reticulum (rER) (fig. S1A). In order to generate a *Cdc50a* conditional allele, we first crossed *Cdc50a* knockout-first allele with *Act-FLPe* mice to remove *FRT* sites. The *Cdc50a* conditional allele was subsequently crossed with *Thy1-CreERT2, -EYFP* mice to generate an inducible neuron-specific *Cdc50a* deletion allele (Fig. 1A). Since homozygous mice for *Cdc50a* knockout-first allele show developmental lethality (*11*), we injected 4-hydroxytamoxifen (4-OHT) or oil into one-month-old *Cdc50a* deletion homozygous allele mice to generate *Cdc50a* conditional knockout (cKO) and control mice, respectively. Compared to control mice, we found that *Cdc50a* cKO mice showed rapid lethality within 2 weeks after 4-OHT injection, without significant changes in body weight or brain morphology (Fig. 1C and fig. S1, B and C). Interestingly, brains from *Cdc50a* cKO and control mice showed no difference in the expression of cleaved caspase 3 (cCas3) and Terminal deoxynucleotidyl transferase dUTP nick end labeling (TUNEL), as well as in the number of NeuN-positive neuronal cells (Fig. 1D and fig. S1, D to F). We also examined processes of *Thy1-CreERT2, -EYFP* positive motor neurons using diaphragm muscles, however, the morphology and number of axons and neuromuscular junctions were all intact in *Cdc50a* cKO compared to control mice (fig. S1G). These data indicate that acute deletion of neuronal CDC50A induces rapid lethality without inducing apparent cell death.

**Fig. 1.**
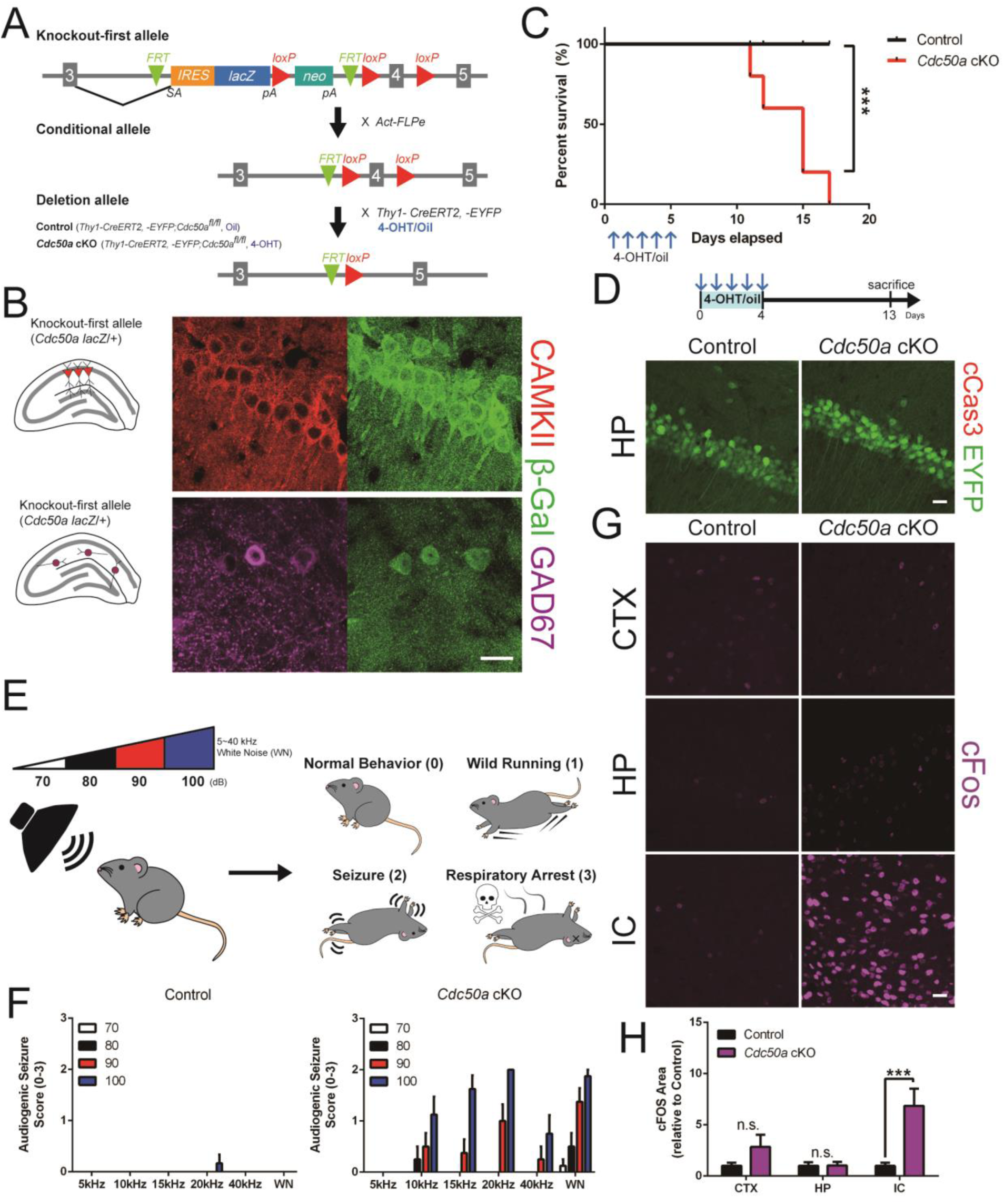
Neuronal Cdc50a deletion induces rapid lethality with audiogenic seizure. (A) Schematic of control and Cdc50a cKO mice generation. (B) Representative confocal images of Cdc50a knockout-first brains immunostained for β-galactosidase (green) with CAMKII (red, upper) or GAD67 (magenta, lower). (C) Survival rates of control and Cdc50a cKO after oil or 4-OHT injection. (D) Representative confocal images of control and Cdc50a cKO brains expressing Thy1-cre/ERT2, -EYFP (green) immunostained for cCas3 (red). (E) Schematic of a behavior test for audiogenic seizure (0: normal behavior, 1: wild running, 2: seizure, and 3: respiratory arrest). (F) Bar graphs showing audiogenic seizure behaviors in control and Cdc50a cKO mice. (G) Representative confocal images of control and Cdc50a cKO brains immunostained for cFos (magenta); CTX, cortex; HP, hippocampus; IC, inferior colliculus. (H) Bar graphs showing the area of cFos expression in control and Cdc50a cKO brains. n = 5 to 8 per group. Bars and error bars indicate mean and SEM, respectively. n.s. = P > 0.05, *P < 0.05, or ***P < 0.001 using Log-rank (Mantel-Cox) test (C) and t-test (G). Scale bar; 20μm (B, D, and G).

Unexpectedly, while monitoring the behaviors of *Cdc50a* cKO mice, we found that *Cdc50a* cKO mice showed convulsive seizures upon exposure to a loud noise (movie S1). To test whether *Cdc50a* cKO mice develop audiogenic seizure, we provided auditory stimuli and scored audiogenic seizure behaviors (0: normal behavior, 1: wild running, 2: seizure, 3: respiratory arrest, movie S2). Interestingly, we found that *Cdc50a* cKO mice showed severe audiogenic seizure behaviors (wild running and seizure) at high frequency between 10 and 20 kHz, compared to control mice (Fig. 1, E and F). In agreement with the audiogenic seizure phenotype, neural activity in *Cdc50a* cKO brains, measured by cFos protein expression at 13 days after 4-OHT injection, was significantly upregulated in the inferior colliculus (IC) region (Fig. 1, G and H), the principal midbrain nucleus of the auditory pathway (*12*).

Next, to test whether audiogenic seizure in *Cdc50a* cKO mice is due to an imbalance between excitatory and inhibitory synapses (*13*), we measured the number of excitatory and inhibitory synapses in various brain regions. Interestingly, there was no significant difference in the number of excitatory synapses, measured by synaptic puncta analysis after both PSD95/VGLUT1 and PSD95/VGLUT2 antibody staining, as well as by sparse labeling of neurons with the tdTomato reporter (Fig. 2, A and B and fig. S2, B to D). Moreover, there was no significant change in the number of inhibitory pre-synapses, by VGAT antibody staining (Fig. 2, C and D). However, surprisingly, we found that the number of inhibitory post-synapses measured by Gephyrin antibody staining, was significantly reduced in *Cdc50a* cKO compared to control mice in the cortex, hippocampus, and IC. Due to the reduced number of Gephyrin positive inhibitory post-synapses, the total number of inhibitory synapses containing both Gephyrin and VGAT was also significantly reduced (Fig. 2, C and D and fig. S2A). Gephyrin is a critical inhibitory post-synaptic element that binds and stabilizes both glycine and γ-aminobutyric acid type A (GABAA) receptors for inhibitory synaptic transmission (*14*). We found that both glycineric and GABAergic post-synapses, measured by glycine receptor and GABAA receptor γ subunit antibodies, respectively, were highly downregulated in *Cdc50a* cKO mice (Fig. 2, F and G and fig. S2A). However, GABAA receptor α and β subunits, which are also localized in the extra-synaptic membrane (*15*), appear to be intact in *Cdc50a* cKO mice (Fig. 2, E and G and fig. S2A), suggesting that *Cdc50a* deletion in neurons specifically reduces the number of inhibitory post-synapses without disrupting general neuronal membranes.

**Fig. 2.**
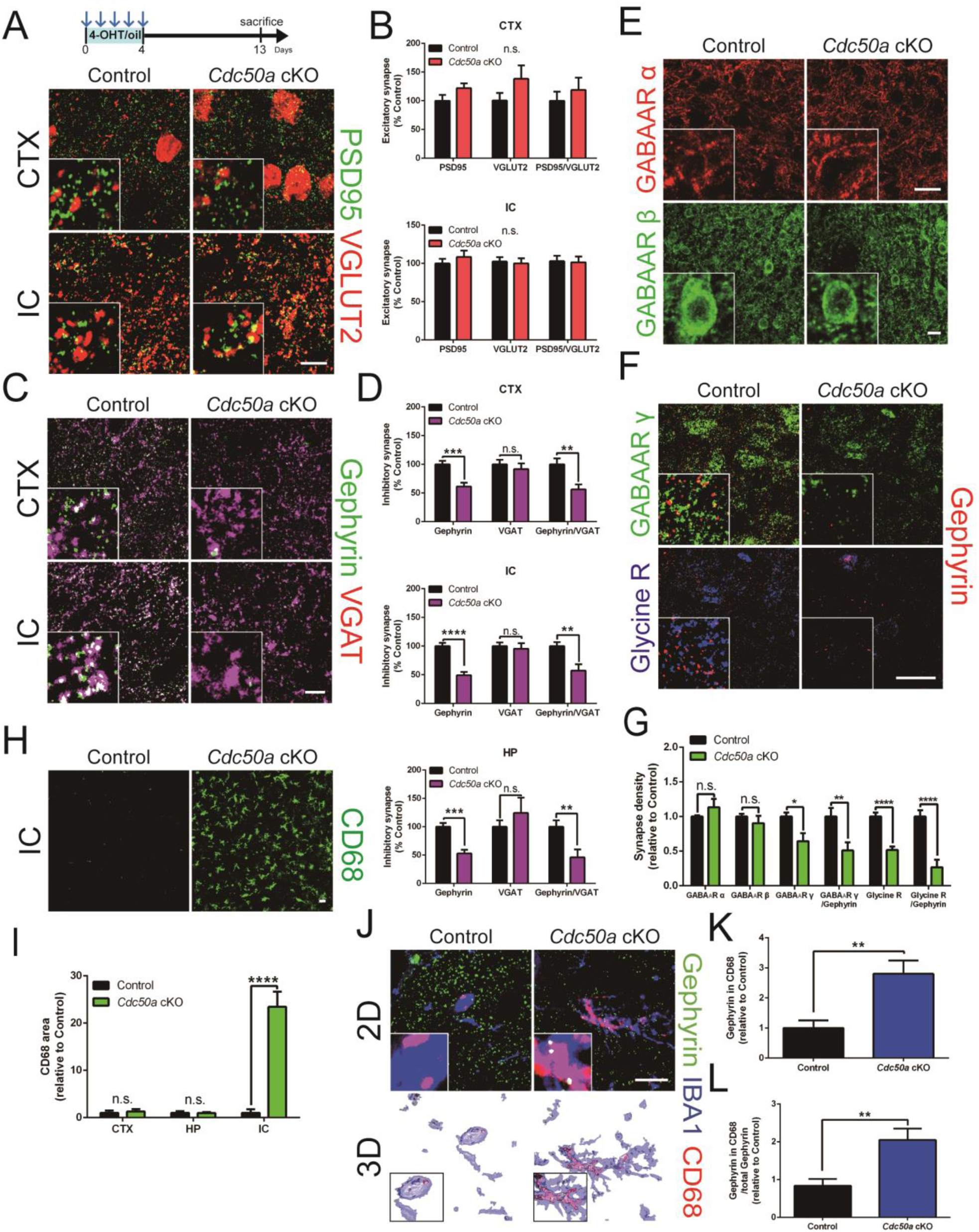
Neuronal *Cdc50a* deletion induces the loss of inhibitory post-synapses by microglial phagocytosis. (A and B) Representative confocal images (A) and bar graphs (B) of excitatory synapses, PSD95 (green) and VGLUT2 (red), in control and *Cdc50a* cKO brains. (C and D) Representative confocal images (C) and bar graphs (D) of inhibitory synapses, Gephyrin (green) and VGAT (magenta), in control and *Cdc50a* cKO brains. (E and F) Representative confocal images of control and *Cdc50a* cKO brains (IC) immunostained for GABAA receptor α (E, red, upper) and β (E, green, lower), and Gephyrin (F, red) with GABAA receptor γ (F, green, upper), and Glycine receptor (F, blue, lower). (G) Bar graphs showing inhibitory post-synapse density in control and *Cdc50a* cKO brains (IC). (H and I) Representative confocal images (H) and bar graph (I) of CD68 (green) in control and *Cdc50a* cKO brains (IC). (J) Representative confocal and 3D reconstituted images of Gephyrin (green) colocalized with CD68 (red) and IBA1 (blue) in control and *Cdc50a* cKO brains (IC). (K and L) Bar graphs showing the area (K) and number (L) of Gephyrin puncta colocalized with CD68 and IBA1 in control and *Cdc50a* cKO brains (IC). Bars and error bars indicate mean and SEM, respectively. *n =* 4 to 6 per group. n.s. = P > 0.05, *P < 0.05, **P < 0.005, ***P < 0.001, or ****P < 0.0001 using *t*-test (C, D, G, I, K, and L). Scale bar; 10μm (A, C, and J) or 20μm (E, F, and H).

Interestingly, when *Cdc50a* cKO mice were housed in a single cage within a soundproof chamber after 4-OHT injection, *Cdc50a* cKO mice were able to survive much longer (fig. S3A). We found that similar to *Cdc50a* cKO mice housed in normal cages (13 days after 4-OHT injection), *Cdc50a* cKO mice housed in a soundproof chamber (20 days after 4-OHT injection) did not show any sign of increased cell death compared to control mice (fig. S3B). However, loss of inhibitory post-synapses in the cortex and IC gradually increased over time after 4-OHT injection (fig. S3C), further indicating that rapid lethality in *Cdc50a* cKO mice results from excessive neuronal hyperactivity due to the loss of inhibitory post-synapses.

To understand why *Cdc50a* cKO mice exhibit an audiogenic seizure phenotype, we compared the extent of PS exposure and reactive gliosis in *Cdc50a* cKO mice housed in normal cages versus soundproof chambers. First, when we dissected out different brain regions from *Cdc50a* cKO mice and used fluorescence-activated cell sorting (FACS) to label PS-positive neurons with AnnexinV, we found that neurons from the IC showed a significant increase in PS exposure at 13 days after 4-OHT injection whereas neurons from the cortex and hippocampus showed significant PS exposure only at 20 days after 4-OHT injection (fig. S3D). Second, we found that in *Cdc50a* cKO mice housed in normal cages, both astrocytes and microglia were highly reactivated in the IC, but not in the cortex and hippocampus. However, the extent of reactive gliosis was significantly increased in all brain regions (IC, cortex, and hippocampus) in *Cdc50a* cKO mice housed in soundproof chambers (fig. S3, E and F). Consistent with the gradual increase of reactive gliosis, the number of cFos-positive cells in the cortex and hippocampus was significantly increased in *Cdc50a* cKO mice housed in a soundproof chamber (fig. S3, G and H). The initial appearance of PS exposure and reactive gliosis in the IC is likely due to strong *Thy1* promoter activity in IC neurons and may explain why *Cdc50a* cKO mice initially show audiogenic seizure phenotype. Indeed, when we sparsely labeled *Thy1-CreERT2, -EYFP* positive neurons with a *Rosa26-CAG-tdTomato* reporter by injecting low doses of tamoxifen, we found that the IC showed the most frequent genetic recombination compared to other brain regions, such as the cortex and hippocampus (fig. S, I and J). Taken together, our data suggest that the IC is the first and primary region that exhibits inhibitory post-synapse loss, reactive gliosis, and excessive excitability in our model. However, these phenotypes spread to other brain regions when *Cdc50a* cKO mice are kept alive for more than 20 days.

Then, how does *Cdc50a* deletion induce loss of inhibitory-post synapses? We hypothesized that microglia and/or astrocytes might phagocytose inhibitory post-synapses by recognizing outer membrane-exposed PS due to *Cdc50a* deletion. To test our hypothesis, we first measured the amount of lysosomes in glial cells and found that total lysosome content, as measured by Cathepsin D antibody staining and microglial lysosome content, as measured by CD68 antibody staining, were both dramatically upregulated in *Cdc50a* cKO mice (Fig. 2, H and I and fig. S4, A and B), suggesting that microglia are responsible for excessive inhibitory post-synapse loss in *Cdc50a* cKO mice. Indeed, we found that Gephyrin-positive puncta, measured by Gephyrin antibody staining (Fig. 2, J to L) and by expression of mCherry-Gephyrin fusion protein (fig. S4, E and F) was highly localized inside CD68 positive microglial lysosomes in *Cdc50a* cKO mice. Since astrocytes also play critical roles in developmental synapse elimination, we also examined whether astrocytes also phagocytose Gephyrin in *Cdc50a* cKO mice. Although we found that the number of LAMP2-positive lysosomes in astrocytes was increased in the cortex, hippocampus, and IC (fig. S4, C and D), mCherry-Gephyrin was mainly localized in microglia rather than astrocytes (fig. S4F), indicating that microglia are mainly responsible for phagocytosing inhibitory-post synapses in *Cdc50a* cKO mice.

*Cdc50a* deficient lymphoma cells stably expose PS throughout the whole outer plasma membrane (*9*). However, since neurons consist of the soma, dendrites and axons, we hypothesized that *Cdc50a* cKO neurons might expose PS in specific compartments, resulting in preferential elimination of inhibitory post-synapses by microglia. To test our hypothesis, we cultured cortical neurons from *Cdc50a* conditional allele mice and transfected *hSyn-mCherry* or *hSyn-Cre-p2a-dTomato* to generate control and *Cdc50a* cKO neurons, respectively. After deleting *Cdc50a*, we monitored the extent of PS exposure at the outer plasma membranes of neurons by utilizing Polarity Sensitive Indicator of Viability (pSIVA), a probe which binds PS reversibly and emits green fluorescence (*16*). Surprisingly, starting from 8 days after *hSyn-Cre-p2a-dTomato* transfection, PS exposure occurred preferentially in the soma first, then gradually spread to the entire in *Cdc50a* cKO neuron (Fig. 3, A and B). Next, to detect this preferential PS exposure in *Cdc50a* cKO brains, we adopted the secreted AnnexinV (secA5) system (*17*), where neighboring cells, in our case, astrocytes, express and secrete AnnexinV fused with mCherry (Fig. 3C). Injecting AAV9-*GFAP-secA5-mCherry* allowed secA5-mCherry to be expressed and secreted from astrocytes, enabling detection of PS-exposed material in brains.

**Fig. 3.**
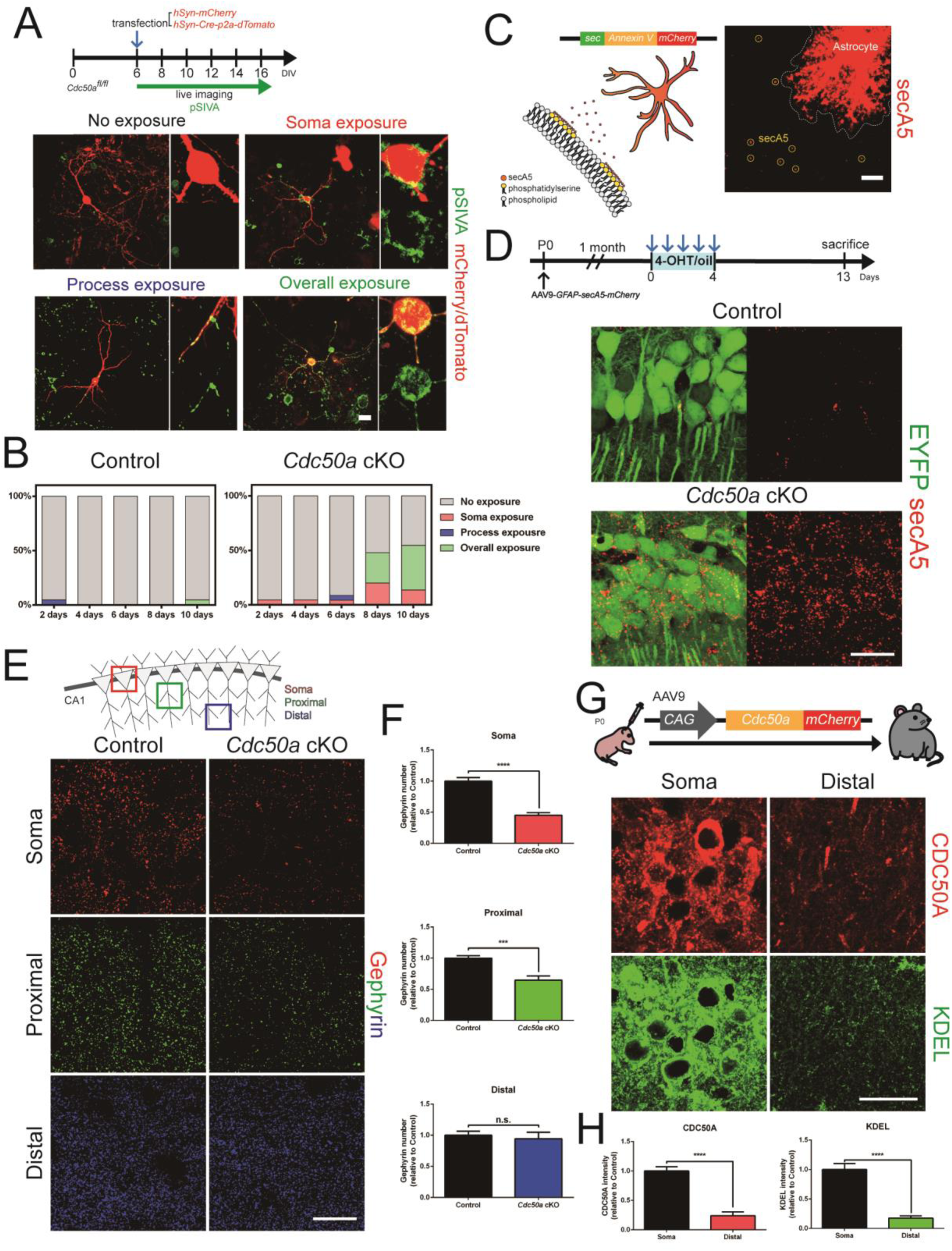
*Cdc50a* cKO induces PS exposure preferentially in neuronal soma. (A) Schematic and representative live confocal images of PS exposure in control and *Cdc50a* cKO neurons *in vitro.* (B) Bar graphs showing percentage of cases with PS exposure in the soma, processes, and overall neurons in control and *Cdc50a* cKO neurons *in vitro*. (C) Schematic and representative confocal images of the secA5 system expressed in astrocytes. (D) Representative confocal images of PS exposure in control and *Cdc50a* cKO (CA1, HP) using AAV9-*GFAP-secA5-mCherry*. (E and F) Representative confocal images (E) and bar graphs (F) of Gephyrin in neuronal soma (red), proximal dendrites (green), and distal dendrites (blue) in control and *Cdc50a* cKO (CA1, HP). (G and H) Representative confocal images (G) and bar graphs (H) of CDC50A*-*mCherry (red) immunostained for KDEL (green) in neuronal soma and distal dendrites in WT (CA1, HP). *n =* 4 to 6 per group (for *in vitro, n =* 20 to 25 per group). n.s. = P > 0.05, *P < 0.05, **P < 0.005, or ****P < 0.0001 using *t-*test (F and H). Scale bar; 10μm (C) or 20μm (A, D, E, and G).

Consistent with our *in vitro* data, we found that secA5 puncta were mostly localized in the neuronal soma rather than in processes in *Cdc50a* cKO brains (Fig. 3, C and D), suggesting that the neuronal soma is the major locus of PS exposure when *Cdc50a* is deleted. Importantly, we also found that the number of Gephyrin-positive inhibitory post-synapses was significantly decreased in the soma and proximal dendrites, but not in distal dendrites of the *Cdc50a* cKO CA1 hippocampus (Fig. 3, E and F). Since CDC50A mainly localizes and functions in the rER, which are abundant in the soma but not in distal dendrites (Fig. 3, G and H), our data suggest that *Cdc50a* cKO preferentially exposes PS in the soma. This may explain the specific loss of inhibitory post-synapses, which are highly concentrated in the neuronal soma, compared to other types of synapses.

Next, to directly address the roles of microglia in the loss of inhibitory post-synapses in *Cdc50a* cKO mice, we adopted two different approaches. First, we depleted microglia in *Cdc50a* cKO brains using PLX3397, a well-characterized colony-stimulating factor 1 receptor (CSF1R) kinase inhibitor, which can deplete microglia efficiently from the brains (*18*). *Cdc50a* cKO and control mice were fed with the same PLX3397 chow for 14 days before injecting 4-OHT or oil, and for an additional 13 days before analysis (Fig. 4A). PLX3397 treatment in control and *Cdc50a* cKO mice mostly depleted microglia in the cortex (fig. S5, B and C), but in the IC, the extent of microglial depletion was variable depending on the individual mouse. However, when we analyzed the correlation between the number of remaining microglia in the IC after PLX3397 treatment (Fig. 4, B and D) and audiogenic seizure score, we found a statistically significant correlation (Fig. 4A and fig. S5A), indicating that microglial ablation was able to rescue the audiogenic seizure phenotype in *Cdc50a* cKO mice. Therefore, we classified PLX3397 treated *Cdc50a* cKO mice into “behavior rescue” and “behavior no-rescue” groups. Next, to test whether microglia ablation can rescue the loss of inhibitory post-synapses in *Cdc50a* cKO mice, we measured the number of inhibitory synapses in five different groups (Control;DMSO group, Control;PLX3397 group, *Cdc50a* cKO;DMSO group, *Cdc50a* cKO;PLX3397 behavior rescue group, and *Cdc50a* cKO;PLX3397 behavior no-rescue group). Interestingly, the *Cdc50a* cKO;PLX3397 behavior rescue group showed a significant rescue in the number of Gephyrin-positive inhibitory-post synapses compared to the *Cdc50a* cKO;DMSO group, whereas the *Cdc50a* cKO;PLX3397 behavior no-rescue group did not (Fig. 4, C and E). These data indicate that microglial depletion is sufficient to rescue the excessive loss of inhibitory synapses and audiogenic seizure in *Cdc50a* cKO mice.

**Fig. 4.**
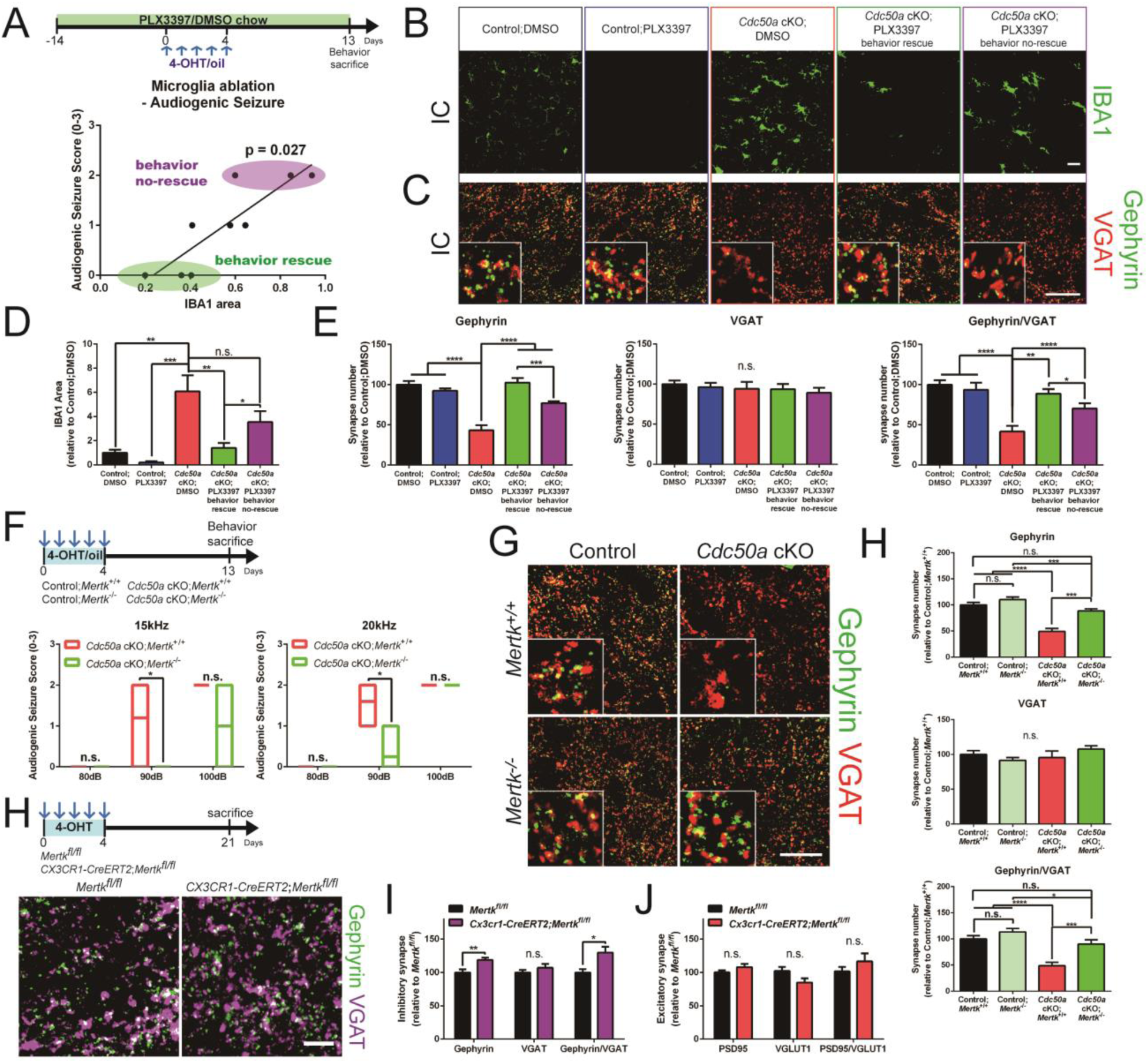
Microglial ablation or *Mertk* deletion rescues the loss of inhibitory post-synapses and audiogenic seizure in *Cdc50a* cKO mice. (A) Bar graphs showing correlation between IBA1 area (IC) and audiogenic seizure score in *Cdc50a* cKO;PLX3397 treated group. (B to E) Representative confocal images of IBA1 (B, green) and Gephyrin(C, green)/VGAT(C, red), and bar graphs showing IBA1 area (D) and inhibitory synapse numbers (E) in control;DMSO, control;PLX3397, *Cdc50a* cKO;DMSO, *Cdc50a* cKO;PLX3397 behavior rescue, and *Cdc50a* cKO;PLX3397 behavior no-rescue groups. (F) Bar graphs showing audiogenic seizure score in *Cdc50a* cKO;*Mertk*^+/+^ and *Cdc50a* cKO;*Mertk*^-/-^ mice. (G and H) Representative confocal images of Gephyrin (G, green) and VGAT (G, red), and bar graphs showing inhibitory synapse numbers (H) in control;*Mertk*^+/+^, *Cdc50a* cKO;*Mertk*^+/+^, control;*Mertk*^-/-^, and *Cdc50a* cKO;*Mertk*^-/-^ brains (IC). (I) Representative confocal images of Gephyrin (green) and VGAT (magenta) in *Mertk*^*fl/fl*^ and *CX3CR1-CreERT2;Mertk*^*fl/fl*^ brains (IC). (J and K) Bar graphs showing the number of inhibitory synapses (J) and excitatory synapses (K) in *Mertk*^*fl/fl*^ and *CX3CR1-CreERT2;Mertk*^*fl/fl*^ brains (IC). *n =* 3 to 9 per group. Bars and error bars indicate mean and SEM, respectively. n.s. = P > 0.05, *P < 0.05, **P < 0.005, ***P < 0.001 or ****P < 0.0001 using ozone correlations test (A), Dunnet’s post hoc test (D, E, and H) and *t-*test (F, J, and K). Scale bar; 20μm (B, C, and G) or 10μm (I).

As a second approach, we deleted critical phagocytic receptors in microglia in the *Cdc50a* cKO background. Previous studies showed that microglia phagocytose excitatory as well as inhibitory synapses through the classical complement pathway during development and neurological diseases (*3-7*). Thus, we crossed complement 3 (*C3*) KO with *Cdc50a* cKO mice to address whether microglia utilize the complement pathway to phagocytose inhibitory-post synapses. Interestingly, although expression of C1q, an initiator for the complement pathway, was significantly enhanced in *Cdc50a* cKO brains (fig. S6A), we found that *C3* KO in the *Cdc50a* cKO background failed to rescue audiogenic seizure, loss of inhibitory synapses, and reactive gliosis compared to the *C3* WT in the *Cdc50a* cKO background (fig. S6, B to F). These data indicate that the complement pathway is not directly associated with regulating the number of inhibitory synapses in *Cdc50a* cKO mice. Apart from the complement pathway, microglia express the MERTK phagocytic receptor (*19*), a member of the TAM (*Tyro3, Axl*, and *Mertk*) tyrosine kinase receptor family, which can recognize PS as an “eat-me” signal through bridging molecules and initiate phagocytosis (*20*). We found that expression of microglial MERTK was highly upregulated in *Cdc50a* cKO mice (fig. S7, A and B). Moreover, to our surprise, introducing a *Mertk* KO in the *Cdc50a* cKO background could mostly rescue audiogenic seizure as well as the loss of inhibitory post-synapses compared to *Mertk* WT in the *Cdc50a* cKO background (Fig. 4, F to H and fig. S7C). However, reactive gliosis still occurred in *Cdc50a* cKO;*Mertk* KO brains, suggesting other phagocytic receptors than *Mertk* might participate in eliminating inhibitory synapses in *Cdc50a* cKO mice (fig. S7D). Taken together, our data strongly suggest that *Cdc50a* cKO mediates the loss of inhibitory synapses through MERTK-dependent microglial phagocytosis.

Next, we investigated whether inhibitory synapses in physiologically normal brains also use PS as a signal for microglia-dependent phagocytosis and synapse elimination. By utilizing our *in vivo* PS reporter system with *secA5-mCherry*, we found that inhibitory post-synapses showed a significantly higher chance of PS exposure than inhibitory pre-synapses (fig. S8, A and B). Interestingly, when we analyzed the PS reporter injected mice at one (juvenile) and three (adult) month-old stages, we found that inhibitory post-synapses no longer showed preferential PS exposure in the adult brains (fig. S8B). In order to label inhibitory synapses and measure their engulfment, we injected AAV9-*hSyn-mCherry-Gephyrin* and AAV9-*GAD67-Synaptophysin-mCherry* for inhibitory post- and pre-synapses, respectively (fig. S7, C to E). Consistent with our PS exposure data, we found that inhibitory post-synapses, but not pre-synapses were frequently found inside Iba1-positive microglia in juvenile mice brains. In agreement with age-dependent changes in PS exposure, the preferential engulfment of inhibitory post-synapses by microglia was not observed in adult brains, suggesting there are age-dependent changes in the extent of inhibitory synapse elimination (fig. S7, C to E). Finally, we deleted *Mertk* specifically in microglia by crossing a *Mertk* conditional allele with *CX3CR1-CreERT2* mice and injecting 4-OHT in one-month-old mice. Surprisingly, we found that deleting *Mertk* in juvenile microglia is sufficient to increase the number of Gephyrin-positive inhibitory post-synapses (Fig. 4, I to K and fig. S7F). Taken together, our data show that PS might serve as a general “eat-me” signal for inhibitory post-synapse elimination in normal juvenile brains through MERTK-dependent microglial phagocytosis.

Since several phagocytic receptors and their bridging molecules recognize PS, it has been proposed that PS may function as an “eat-me” signal for synapse elimination (*20*). However, *in vivo* genetic evidence to address such claims have been lacking. By deleting an upstream modulator of PS in mature neurons, we reveal that CDC50A-dependent PS exposure is specifically required for eliminating inhibitory post-synapses by microglia. This unexpected finding raises many intriguing questions. How does PS exposure in the soma induced by CDC50A deletion specifically initiate inhibitory post-synapse elimination without damaging extra-synaptic membranes? Are other upstream modulators of PS, such as scramblases involved in synapse elimination as well? How does neuronal activity interplay with PS exposure at synapses? Interestingly, excitatory-inhibitory imbalance has been reported in many neurological disorders, such as autism spectrum disorder, schizophrenia, frontotemporal dementia, and several forms of seizures (*7, 21*). Our finding strongly suggests that such imbalance can originate from abnormal PS exposure at synapses or hyperactivated MERTK-dependent microglia phagocytosis. Therefore, modulating inhibitory synapse elimination by microglia may serve as a novel strategy for treating various brain disorders.

## Materials and Methods

### Animals

*Cdc50a*^tm1a(KOMP)Wtsi^ mice were obtained from KOMP and crossed with B6;SJL-Tg(*ACT-FLPe*)9205Dym/J mice, gifted from Dr. Jeong Ho Lee (KAIST), to remove the *FRT* sites. In order to generate *Cdc50a* deletion allele, *FRT* deleted mice were bred with Tg(*Thy1-CreERT2,-EYFP*)HGfng/PyngJ mice (Jackson Laboratory). *Thy1-CreERT2,-EYFP*;*Cdc50a*^*fl/fl*^ mice were used for control and *Cdc50a* cKO mice. Control and *Cdc50a* cKO mice were intraperitoneally injected with oil or 4-OHT (Sigma, 75μg per body weight (g)), respectively. One-month-old mice were injected with oil or 4-OHT once a day for consecutive 5 days and sacrificed after 13 days or 20 days. To sparsely label *Cdc50a* deleted cells, *Thy1-CreERT2,-EYFP*;*Cdc50a*^*fl/fl*^ mice were crossed with B6.Cg-*Gt(ROSA)26Sor*^*tm3(CAG-EYFP)Hze*^/J (Jackson Laboratory). *Thy1-CreERT2,-EYFP*; *Cdc50a*^*fl/fl*^; *Gt(ROSA)26Sor*^*tm3(CAG-EYFP)Hze*^/J mice were intraperitoneally injected once with tamoxifen (Sigma, 7.5μg per body weight (g)) in one-month-old ages and sacrificed after 24 days. In order to knock-out phagocytic receptors in control and *Cdc50a* cKO mice, B6;129S4-*C3*^*tm1Crr*^/J and B6;129-*Mertk*^*tm1Grl*^*/J* mice (Jackson Laboratory) were crossed with *Thy1-CreERT2,-EYFP*;*Cdc50a*^*fl/fl*^ mice, respectively. For microglial *Mertk* KO, *Mertk*^*fl/fl*^ mice were crossed with B6.129P2(Cg)-*Cx3cr1*^*tm2.1(cre/ERT2)Litt*^/WganJ mice, gifted from Dr. Seyun Kim (KAIST). *Mertk*^*fl/fl*^ and B6.129P2(Cg)-*Cx3cr1*^*tm2.1(cre/ERT2)Litt*^/WganJ;*Mertk*^*fl/fl*^ were intraperitoneally injected with 4-OHT (Sigma, 75μg per body weight (g)) for control and *Mertk* cKO, respectively. *Mertk* floxed mouse line was generated using clustered regularly interspaced short palindromic repeats (CRISPR) technology (Applied StemCell). Briefly, a mixture of two sets of active guide RNA molecules (gRNAs), two single stranded oligodeoxylnucleotides (ssODNs) and qualified Cas-9 mRNA were injected into the cytoplasm of C57BL/6 embryo and 2 *LoxP* cassettes was inserted into introns 1 and 2 each to flank the exon 2 of *Mertk* locus. For genotyping, the primers are listed as below: Forward: 5’-CTTCATCATGCTCACCTCAAACC-3’, Reverse: 5’-GTGCAGAATATTCACCT GACTGC-3’. Both sexes were included in all experiments. WT C57BL/6N mice were purchased from DBL and Samtaco. All procedures were approved by Institutional Animal Care and Use Committees (IACUC) protocols from Korea Advanced Institute of Science and Technology.

### Immunohistochemistry and Image Analysis

Animals were anesthetized by isofluorane (Piramal) or avertin (2, 2, 2-Tribomoethanol, Sigma) and transcardially perfused by phosphate buffered saline (PBS) followed by 4% paraformaldehyde (PFA, WAKO). Brains and diaphragm were dissected out, and post-fixed for overnight in 4% PFA at 4°C, and then cryoprotected in 30% sucrose (Sigma)/PBS for 24 hours at 4°C. After embedding and freezing brain samples in OCT compound (Leica), serial brain sections (30μm) were collected in 24 well plates filled with PBS. Then, each sections were treated with blocking solution (4% bovine serum albumin, 0.3% triton X-100 in PBS) at room temperature (RT) for one hour, and incubated with the following primary antibodies in the blocking solution; anti-CAMKII (rabbit monoclonal from Abcam), anti-β-galactosidase (chicken polyclonal from Aves Labs), anti-GAD67 (mouse monoclonal from Merck), anti-cleaved Caspase-3 (rabbit polyclonal from Cell Signaling Technology), anti-NeuN (mouse monoclonal from Merck), anti-cFos (rabbit polyclonal from Cell Signaling Technology), anti-mCherry (rat monoclonal from Invitrogen), anti-KDEL (rabbit monoclonal from Abcam), anti-PSD95 (rabbit polyclonal from Invitrogen), anti-VGLUT1 (guinea pig polyclonal from Merck), anti-VGLUT2 (guinea pig polyclonal from Merck), anti-Gephyrin (mouse monoclonal from SYSY and rabbit chimeric monoclonal from SYSY), anti-VGAT (rabbit polyclonal from SYSY and guinea pig polyclonal from SYSY), anti-GABA A Receptor α1 (rabbit polyclonal from Alomone labs), anti-GABA A Receptor β 2,3 (mouse monoclonal from Merck), anti-GABA A Receptor γ (guinea pig polyclonal from SYSY), Glycine receptor (mouse monoclonal from SYSY), anti-IBA1 (rabbit polyclonal from Wako, goat polyclonal from Novus Biologicals), anti-GFAP (mouse monoclonal from Merck), CD68 (rat monoclonal from Abcam), anti-Cathepsin D (goat polyclonal from R&D systems), anti-LAMP2 (rat monoclonal from Abcam), anti-S100β (rabbit monoclonal from Abcam), anti-MERTK (rat monoclonal from Invitrogen), anti-C1q (rabbit monoclonal from Abcam). Sections were rinsed 5 times with PBST (0.1% Tween 20 in PBS, Sigma), and incubated with Alexa fluorophore secondary antibodies (Invitrogen) overnight at 4°C. After rinsing with PBST serial times, sections were coverslipped with antifade mounting medium with DAPI or without DAPI (Vectorshield). For anti-GABA A Receptor γ and Glycine receptor antibody staining, brain sections were additionally treated with citrate antigen retrieval and methanol/acetic acid before blocking, respectively. For TUNEL assay, brain sections were treated with in situ Cell Death Detection Kit (Sigma).

All images were acquired with LSM880 confocal microscope (Carl Zeiss). All images were processed using Image J (Fiji) and Imaris software for 3D reconstruction. For colocalization and synapse number analysis, data were processed with ImageJ Distance Analysis (DiAna) plugin (*22*).

### Plasmid and Virus preparation

To generate AAV-*CAG-Cdc50a-mCherry, Mus musculus Cdc50a* cDNA and *mCherry* (Addgene) sequences were amplified by overlap PCR and subcloned into the AAV-*CAG-GFP* (Addgene) at BamHI and EcoRI. For *Cdc50a* cDNA, total RNA was extracted from C57BL/6N mice brain using Rneasy Plus kit (Qiagen) and performed cDNA synthesis using reverse transcriptase (Promega). The primers for *Cdc50a* cDNA are listed as below: Forward: 5’-ATGGCGATGAACTATAGCGCGAA-3’, Reverse: 5’-AATGGTGATGTCAGCAGTGT-3’.

To generate AAV-*hSyn-mCherry-Gephyrin, Gephyrin* sequences were amplified by PCR from *pmCherryC2-Gephyrin P1* (Addgene) and subcloned into the pAAV-*hSyn-mCherry* (Addgene) at BsrGI. AAV-*GAD67-synaptophysin-mCherry-GFP* was obtained from our unpublished study.

To generate AAV-*CAG-secA5-YFP, secA5-YFP* sequences were amplified by PCR from pBH-*UAS-secA5-YFP* (Addgene), and subcloned into the pAAV-*CAG-GFP* (Addgene) at EcoRI and SalI. For AAV-*GFAP-secA5-mCherry, secA5* sequences were amplified by PCR from pBH-*UAS-secA5-YFP* (Addgene) and *mCherry* sequences were amplified by PCR from pAAV-*hSyn-mCherry* (Addgene) sequences. *secA5-mCherry* sequences were amplified by overlap PCR from *secA5* and *mCherry*, and subcloned into the pAAV-*GFAP-EGFP* (Addgene) at SalI and EcoRI.

For *in vitro* studies, pAAV-*hSyn-mCherry* (Addgene) and *hSyn-Cre-p2a-dTomato* (Addgene) plasmids were used.

AAV preparation was performed by previous described methods with minor modifications (*23*). Briefly, HEK293 cell lines were transfected with transfer plasmid (ITR containing plasmid), helper plasmid (pAdDeltaF6, Addgene), and capsid plasmid (pAAV9, Addgene) by polyethylenimine (PEI, Sigma). After 72 hours, transfected cells were harvested through cell scraper and centrifugation, and lysed with freeze/thawing in hypotonic buffer containing Dulbecco’s Modified Eagle Medium (DMEM, Welgene) and deoxyribonuclease I (DNase I, Worthington). After removing cell debris through centrifugation, chloroform (Sigma) was applied in supernatant, and top aqueous phase was collected through centrifugation. PEG8000 (Sigma) and ammonium sulfate (Sigma) solutions were applied in aqueous phase, and bottom clear phase was collected through centrifugation. Amicon Ultra (Merck) was used for concentration.

### Virus injection

C57BL/6N WT pups (P0) were anesthetized with hypothermia (for no more than four minutes). Under the anesthesia, virus with Trypan Blue (Gibco, 1µl) was injected into the lateral ventricle using glass pipette (WPI). After viral injections, pups were recovered on the heating pad, and returned to their home cages. For AAV9-*CAG-Cdc50a-mCherry* injection, mice were sacrificed after one month. For AAV9-*hSyn-mCherry-Gephyrin*, AAV9-*GAD67-synaptophysin-mCherry* and AAV9-*GFAP-secA5-mCherry* injection, mice were sacrificed after one or three months for the analysis of juvenile and adult stages, respectively. For control and *Cdc50a* cKO experiment, mice pups (P0) were injected AAV9-*hSyn-mCherry-Gephyrin* or AAV9-*GFAP-secA5-mCherry* virus and sacrificed 13 days after oil or 4-OHT injection.

### Primary neuron cultures and pSIVA live imaging

Primary neuron cultures were prepared from *Cdc50a* conditional allele mice pups (P0). Briefly, cerebral cortex was dissected out in Hank’s Buffered Salt Solution (HBSS, Gibco) and digested by trypsin-EDTA (Gibco). Digested cells were further dissociated by gentle trituration with Minimum Essential Media (MEM, Gibco) containing 10% Fetal Bovine Serum (FBS, Gibco), 25mM glucose (Sigma), 2mM glutamine (Gibco) and 1% penicillin/streptomycin (Gibco). Cell suspension was plated on poly-D-lysine and laminin pre-coated glass coverslip (Corning). At two hours after plating, media was changed with Neurobasal A (Gibco) containing 2mM glutamine (Gibco), B27 (Gibco), and 1% penicillin/streptomycin (Gibco). Cells were incubated at 37°C humidified incubator with 5% CO2 in air with media changes twice a week. Cells were transfected by lipofectamine 2000 (ThermoFisher) at 6 days *in vitro* (DIV), and imaged in DIV8, DIV10, DIV12, DIV14, and DIV16. For PS exposure, pSIVA-IANBD (Abcam) was incubated with transfected cells for 10 minutes before live cell imaging. All images were acquired with LSM880 confocal microscope (Carl Zeiss) during live cell status.

### PLX3397 treatment

PLX3397 (pexidartinib) was purchased from SelleckChem, and dissolved in Dimethyl sulfoxide (DMSO) for 50mM stock concentration. PLX3397 was formulated in rodent standard chow (290mg/kg). For vehicle, DMSO was also formulated in rodent standard chow. Control and *Cdc50a* cKO mice were fed with DMSO chow and PLX3397 chow during 4 weeks.

### Audiogenic seizure behavior

Mice were placed in an acrylic chamber (32*31*21.5 cm) and covered with a transparent acrylic lid. Each sound (5kHz, 10kHz, 15kHz, 20kHz, 40kHz and white noise) was presented from speaker for one minute and rested for one minute for four cycles, and sound intensity were serially increased in each cycle (70dB, 80dB, 90dB and 100dB). All auditory stimuli were generated by Matlab software. All behaviors were recorded by a video camera. Mice was scored as four different behavior types (0: normal response, 1: wild running, 2: seizure, and 3: respiratory arrest) during auditory stimulus.

### Western blot

Control and *Cdc50a* cKO mice brains were dissected out and lysed in EzRIPA lysis buffer (Atto) with protease inhibitors (Atto) and phosphatase inhibitor (Atto). After proteins were quantified by the Bradford method (Bio-Rad), lysates were mixed with 5x SDS sample buffer (250mM Tris-HCl pH6.8, 5% 2-Mercaptoethanol, 10% sodium dodecyl sulfate, 0.5% Bromophenol blue, and 50% Glycerol). Samples were loaded on SDS-PAGE gel (Bio-Rad) and transferred to polyvinylidene fluoride (PVDF, Atto) membrane. The membranes were blocked in blocking solution (10% skim milk in Tris buffered saline, TBS) for one hour at room temperature and incubated with following primary antibodies in the blocking solution (5% skim milk in TBS) at 4°C. Homer (mouse monoclonal from SYSY), Synapsin (rabbit monoclonal from Cell Signaling Technology), Gephyrin (mouse monoclonal from SYSY), GABA A Receptor β 2, 3 (mouse monoclonal from Merck), GABA A Receptor γ (guinea pig polyclonal from SYSY), Glycine receptor (mouse monoclonal from SYSY), and GAPDH (rabbit monoclonal from Santa Cruz). Membranes were rinsed 3 times in TBST (0.1% Tween 20 in TBS), and incubated with HRP conjugated secondary antibodies (Santa Cruz and Invitrogen) for one hour at room temperature. After washing with TBST for 3 times, membranes were developed by Enhanced Chemiluminescence (ECL, GE Healthcare), and visualized with Chemidoc XRS+ (Bio-Rad). Data were analyzed with ImageJ (Fiji) software.

### FACS

Control and *Cdc50a* cKO mice were transcardially perfused with PBS, and cortex, hippocampus, and IC were dissected out. Each brain region was chopped and digested by papain (Worthington). Digested tissues were further dissociated by gentle trituration, and centrifuged using 22% percoll (GE Healthcare) to remove myelin debris. After removing supernatant, cells were incubated with AnnexinV-Cy5 Apoptosis Kit (Biovision) at RT for 10 minutes. Samples were analyzed with FACS LSR Fortessa (BD Bioscience) and data were analyzed with Flowjo software.

### Statistical Analysis

GraphPad Prism 6 software was used for all statistical analysis. For comparing two samples, data were tested using unpaired *t*-test or multiple *t*-test with 95% confidence. For comparing multiple samples, analysis of variance (ANOVA) with Dunnet’s post hoc test was used. In order to compare the survival test, survival curves were analyzed with Log-rank (Mantel-Cox) test with 95% confidence. For correlation test, ozone correlations were used. All data were presented as means and standard error of the mean (SEM).

## Acknowledgments

We thank all members of the Chung laboratory for helpful discussion. This work was supported by grants from the Samsung Science & Technology Foundation (BA170110787, W.-S. C.).

## Author contributions

W.-S.C. and J.P. designed projects. J.P. performed all the experiments including plasmid cloning, virus production, cell culture, immunohistochemistry, confocal imaging, western blot, FACS analysis and behavior analysis. E.J. and S.-H.L. provided a software with auditory stimulus. W.-S.C. and J.P. wrote the paper.

## Data and materials availability

All data is available in the main text or the supplementary materials.

**Fig. S1.**
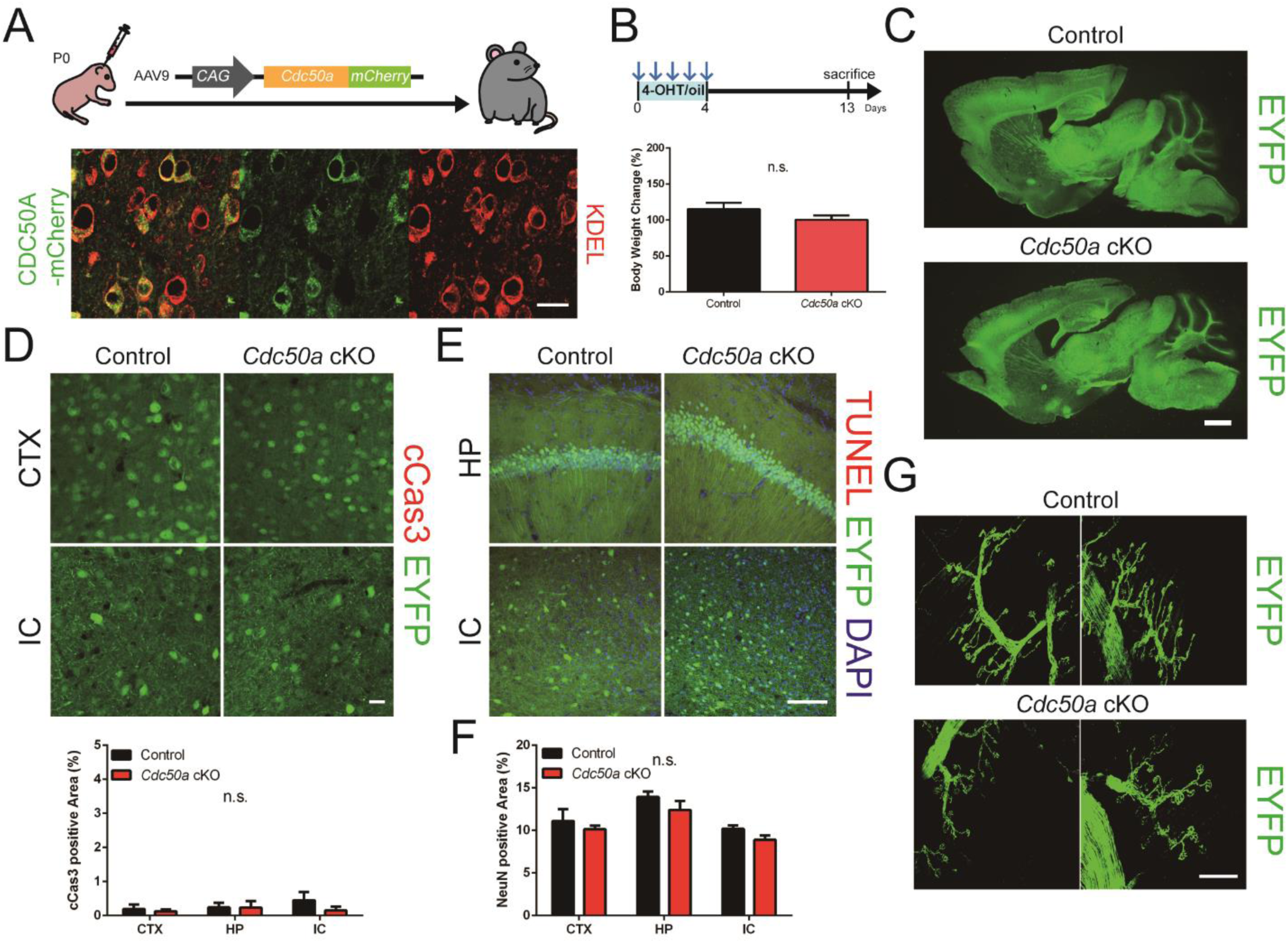
Neuronal *Cdc50a* ablation does not induce apparent cell death. (A) Representative confocal images of CDC50A*-*mCherry (green) immunostained for KDEL (red) in WT brains (CTX). (B) A bar graph showing body weight change in control and *Cdc50a* cKO mice. (C) Representative brain images of EYFP (green) expressing neurons from control and *Cdc50a* cKO mice. (D) Representative confocal images (upper) and bar graphs (lower) of EYFP (green) expressing control and *Cdc50a* cKO brains immunostained for cCas3 (red). (E) Representative confocal images of EYFP (green) expressing control and *Cdc50a* cKO brains stained for TUNEL (red) and DAPI (blue). (F) Bar graphs showing the area of NeuN in the control and *Cdc50a* cKO brains. (G) Representative confocal images of EYFP expressing axons and neuromuscular junctions from motor neurons in the control and *Cdc50a* cKO diaphragms. *n =* 4 to 6 per group. Bars and error bars indicate mean and SEM, respectively. n.s. = P > 0.05 or ****P < 0.0001 using *t-*test (B, D, and F). Scale bar; 20μm (A and D) or 100μm (B, E and G).

**Fig. S2.**
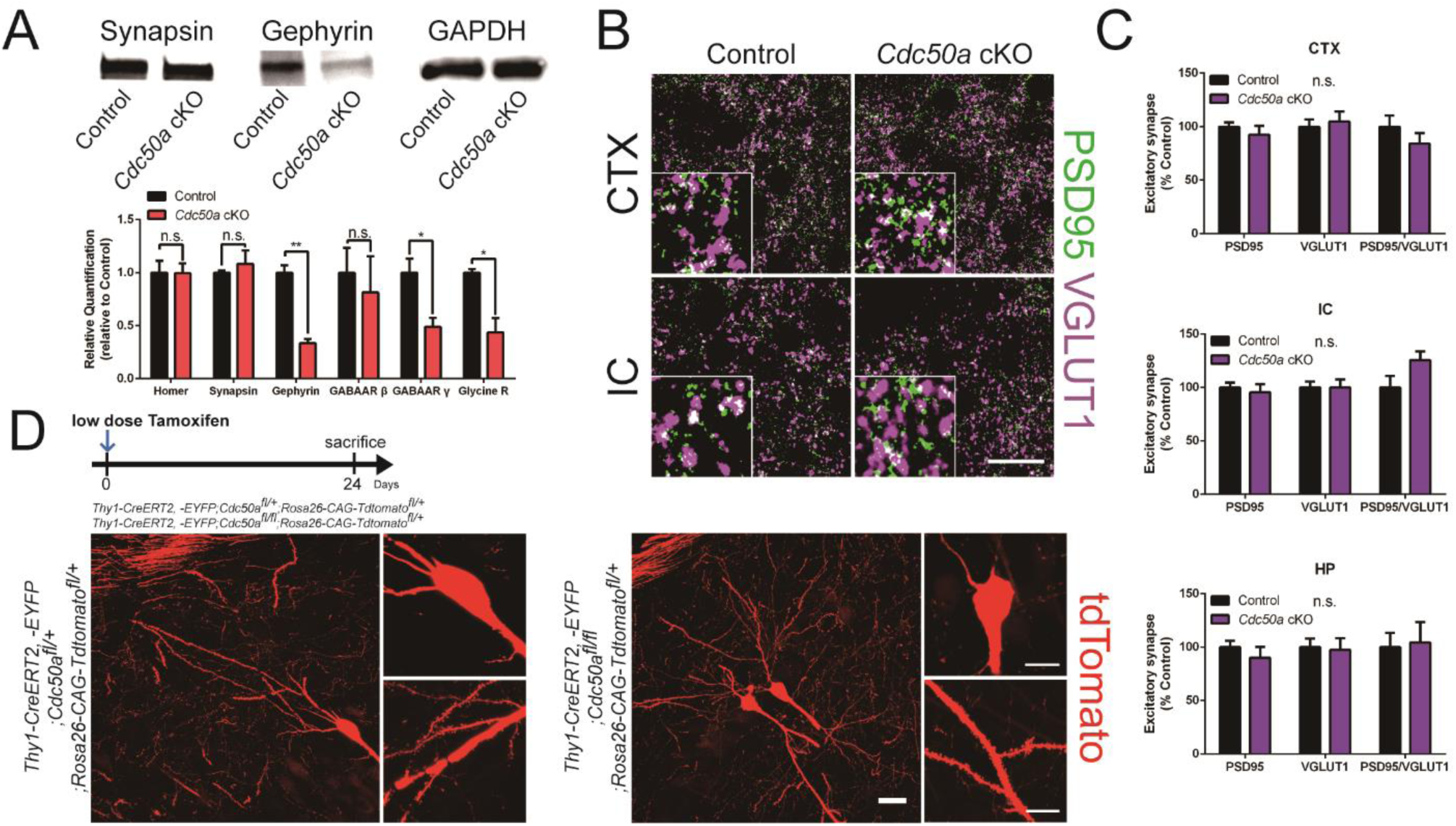
Neuronal *Cdc50a* ablation does not affect excitatory synapses. (A) Representative western blot images showing Synapsin, Gephyrin and GAPDH from control and *Cdc50a* cKO brains (IC, upper). Bar graphs showing relative quantification of western blots with synaptic proteins from control and *Cdc50a* cKO brains (IC, lower). (B and C) Representative confocal images (B) and bar graphs (C) of PSD95 (green) and VGLUT1 (magenta) positive excitatory synapses in the control and *Cdc50a* cKO brains. (D) Representative confocal images of tdTomato (red) expressing neurons from *Thy1-CreERT2, - EYFP*;*Cdc50a*^*fl/+*^;*Rosa26-CAG-tdTomato* ^*fl/+*^ and *Thy1-CreERT2, -EYFP*;*Cdc50a*^*fl/fl*^;*Rosa26-CAG-tdTomato* ^*fl/+*^ brains after low dose tamoxifen injection. Bars and error bars indicate mean and SEM, respectively. *n =* 3 to 6 per group. n.s. = P > 0.05, *P < 0.05 or **P < 0.005 using *t-* test (A and C). Scale bar; 10μm (B and D) or 20μm (D).

**Fig. S3.**
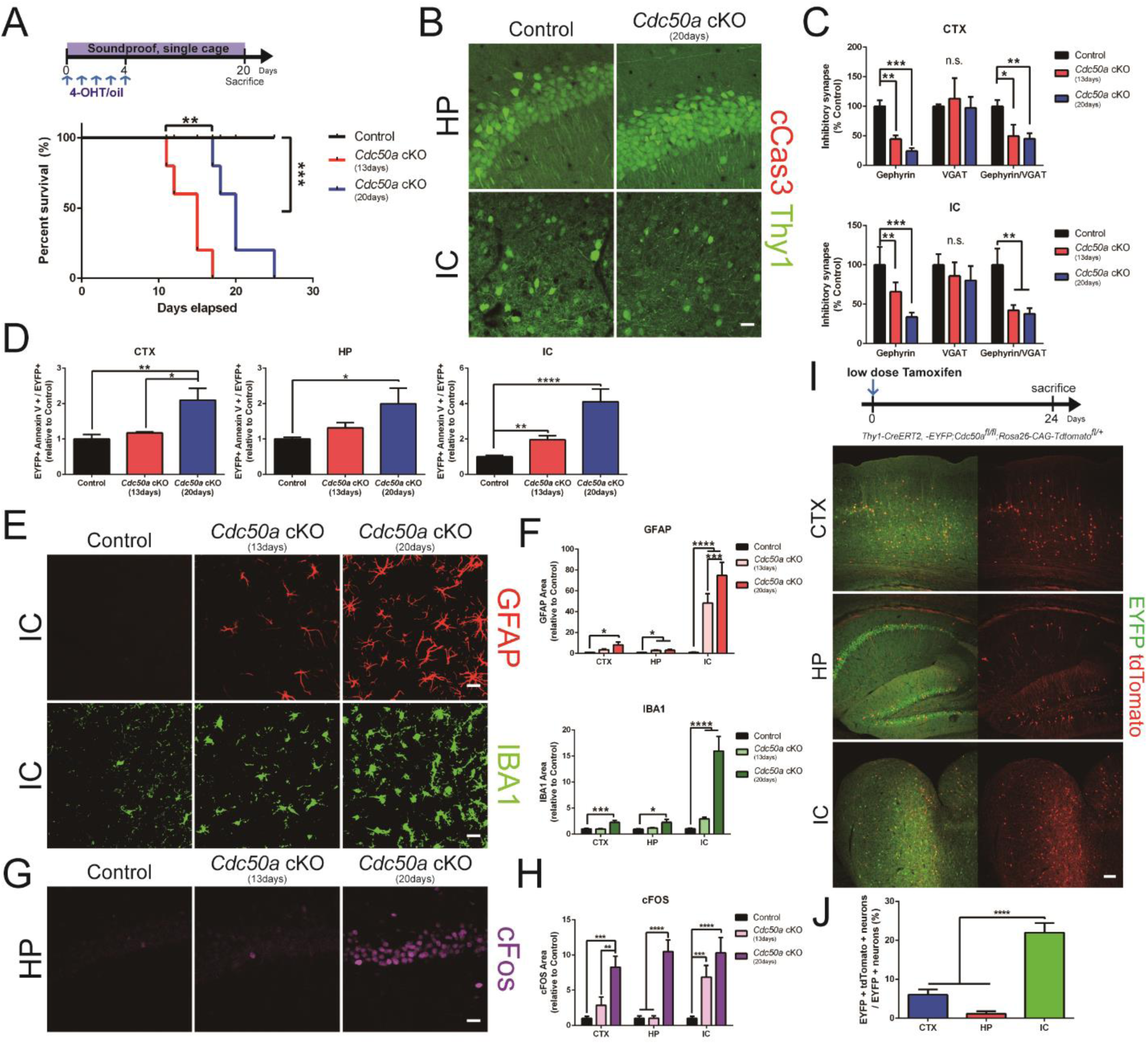
Soundproofing rescues rapid lethality, but not excessive inhibitory synapse loss and reactive gliosis in *Cdc50a* cKO mice. (A) Survival rates of control, *Cdc50a* cKO mice in normal cages, and *Cdc50a* cKO mice housed in a single cage within a soundproof chamber. (B) Representative confocal images of EYFP (green) expressing control and *Cdc50a* cKO (20 days) brains immunostained for cCas3 (red). (C) Bar graphs showing the number of inhibitory synapses of control, *Cdc50a* cKO (13 days), and *Cdc50a* cKO (20 days) brains. (D) Bar graphs showing percentage of EYFP and AnnexinV double positive neurons from CTX, HP, and IC of control, *Cdc50a* cKO (13days), and *Cdc50a* cKO (20days) brains. (E and F) Representative confocal images (E) and bar graphs (F) of GFAP (red) and IBA1 (green) in control, *Cdc50a* cKO (13days), and *Cdc50a* cKO (20days) brains. (G and H) Representative confocal images (G) and bar graphs (H) of cFos (magenta) in control, *Cdc50a* cKO (13days), and *Cdc50a* cKO (20days) brains. (I) Representative confocal images of tdtomato (red) expressing neurons from *Thy1-CreERT2, - EYFP*;*Cdc50a*^*fl/fl*^;*Rosa26-CAG-tdTomato* brains expressing EYFP (green) after low dose Tamoxifen injection. (J) Bar graphs showing percentage of tdTomato positive neurons out of total EYFP expressing neurons in *Thy1-CreERT2, -EYFP*;*Cdc50a*^*fl/fl*^;*Rosa26-CAG-tdTomato* brains. *n =* 4 to 8 per group. Bars and error bars indicate mean and SEM, respectively. n.s. = P > 0.05, *P < 0.05, **P < 0.005, ***P < 0.001 or ****P < 0.0001 using Log-rank (Mantel-Cox) test (A) and Dunnet’s post hoc test (C, D, F, H, and J). Scale bar; 20μm (B, E, and G) or 100μm (J).

**Fig. S4.**
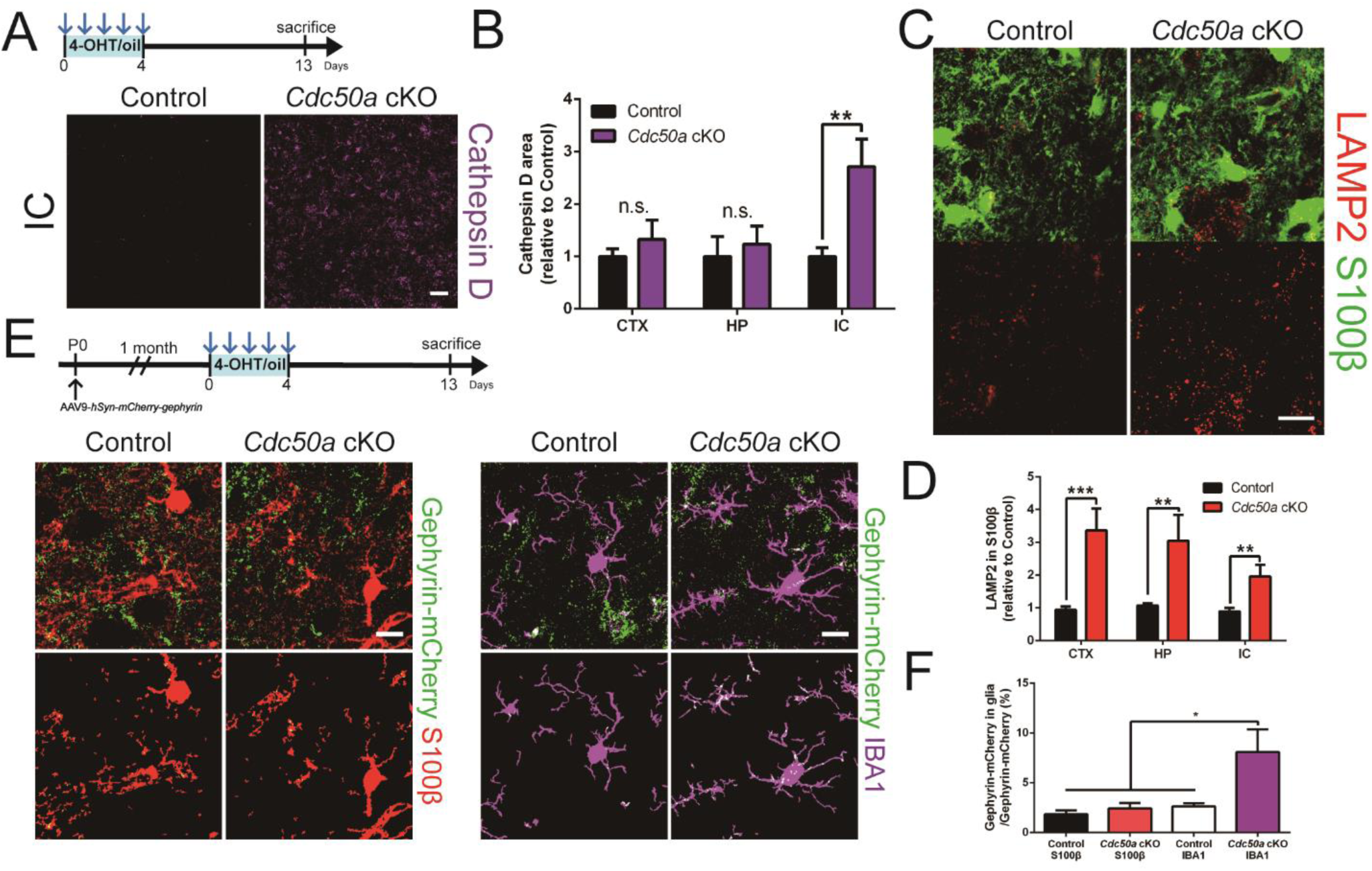
Neuronal *Cdc50a* ablation increases glial lysosome content and microglial inhibitory post-synapse phagocytosis. (A and B) Representative confocal images (A) and bar graphs (B) of Cathepsin D (magenta) in control and *Cdc50a* cKO brains. (C and D) Representative confocal images (C) and bar graphs (D) of LAMP2 (red) and S100β (green) in control and *Cdc50a* cKO brains (IC). (E) Representative confocal images (upper) and colocalization images (lower) of mCherrry-Gephyrin (green) with S100β (red, left) and IBA1 (magenta, right) in control and *Cdc50a* cKO brains (CTX). (F) Bar graphs showing the area of engulfed mCherrry-Gephyrin inside S100β or IBA1-positive cells normalized by the area of total mCherrry-Gephyrin puncta in control and *Cdc50a* cKO brains (CTX). *n =* 4 to 6 per group. Bars and error bars indicate mean and SEM, respectively. n.s. = P > 0.05, *P < 0.05, **P < 0.005, or ***P < 0.001 using *t-*test (B and D) and Dunnet’s post hoc test (F). Scale bar; 10μm (E), 20μm (C), or 40μm (A).

**Fig. S5.**
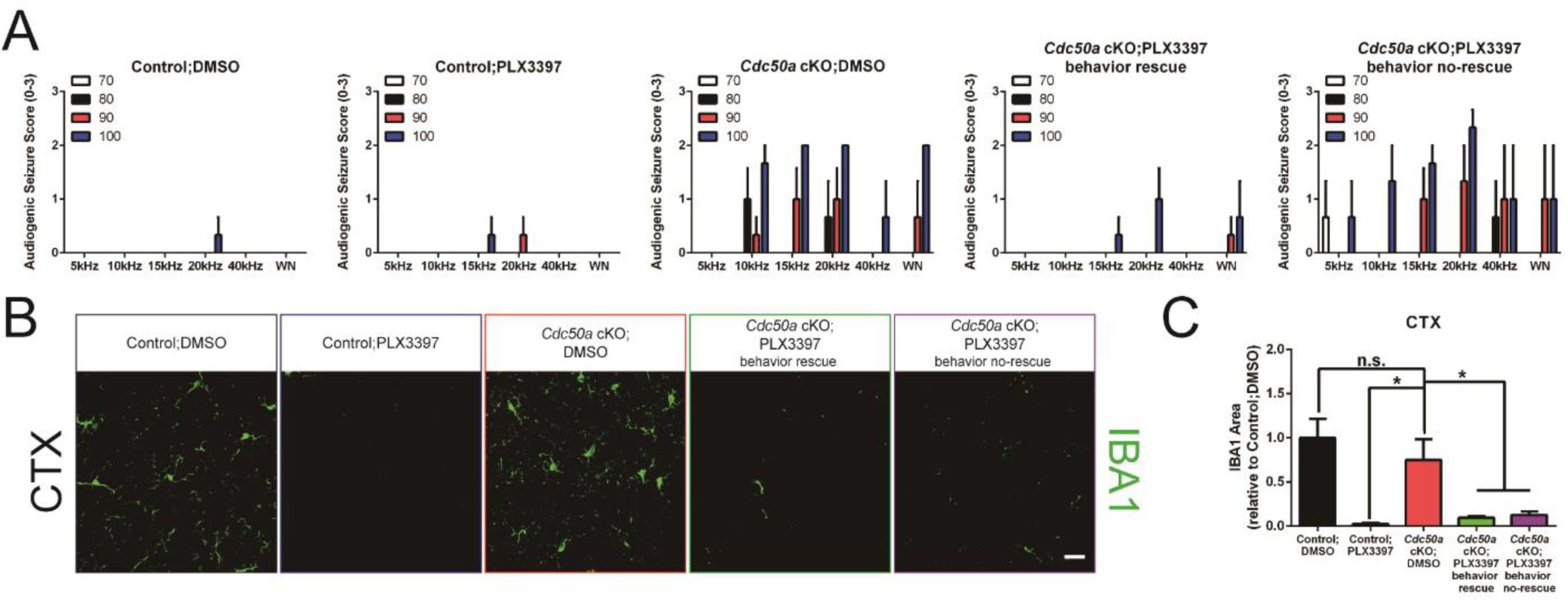
Microglia ablation rescues audiogenic seizure in *Cdc50a* cKO mice. (A) Bar graphs showing audiogenic seizure score in control;DMSO, control;PLX3397, *Cdc50a* cKO;DMSO, *Cdc50a* cKO;PLX3397 behavior rescue, and *Cdc50a* cKO;PLX3397 behavior no-rescue groups. (B and C) Representative confocal images (B) and bar graphs (C) of IBA1 positive cells in control;DMSO, control;PLX3397, *Cdc50a* cKO;DMSO, *Cdc50a* cKO;PLX3397 behavior rescue, and *Cdc50a* cKO;PLX3397 behavior no-rescue groups. *n =* 3 to 6 per group. Bars and error bars indicate mean and SEM, respectively. n.s. = P > 0.05 or *P < 0.05 using Dunnet’s post hoc test (C). Scale bar; 20μm (B).

**Fig. S6.**
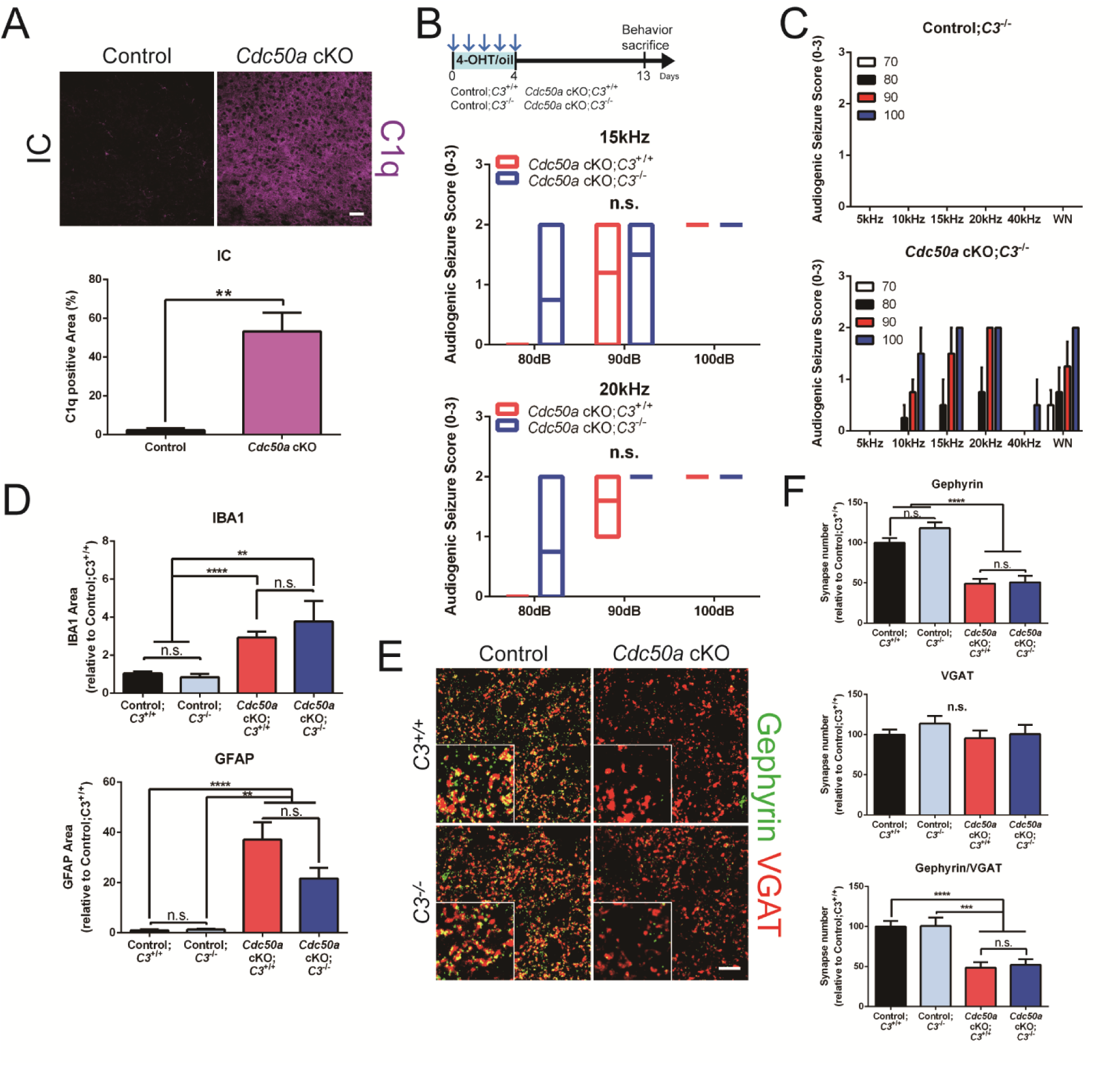
*C3* deficiency fails to rescue the loss of inhibitory synapses and audiogenic seizure in *Cdc50a* cKO mice. (A) Representative confocal images (upper) and bar graphs (lower) of C1q (magenta) in control and *Cdc50a* cKO brains. (B) Bar graphs showing audiogenic seizure score in *Cdc50a* cKO;*C3*^+/+^ and *Cdc50a* cKO;*C3*^-/-^. (C) Bar graphs showing audiogenic seizure score in control;*C3*^-/-^, and *Cdc50a* cKO;*C3*^-/-^ mice. (D) Bar graphs showing the area of IBA1 (upper) and GFAP (lower) positive cells in control;*C3*^+/+^, *Cdc50a* cKO;*C3*^+/+^, control;*C3*^-/-^, and *Cdc50a* cKO;*C3*^-/-^ brains (IC). (E and F) Representative confocal images (E) and bar graphs (F) of Gephyrin (green) and VGAT (red) positive inhibitory synapses in control;*C3*^+/+^, *Cdc50a* cKO;*C3*^+/+^, control;*C3*^-/-^, and *Cdc50a* cKO;*C3*^-/-^ brains (IC). *n =* 4 to 6 per group. Bars and error bars indicate mean and SEM, respectively. n.s. = P > 0.05, **P < 0.005, ***P < 0.001 or ****P < 0.0001 using *t-*test (A and B) and Dunnet’s post hoc test (D and F). Scale bar; 10μm (E) or 40μm (A).

**Fig. S7.**
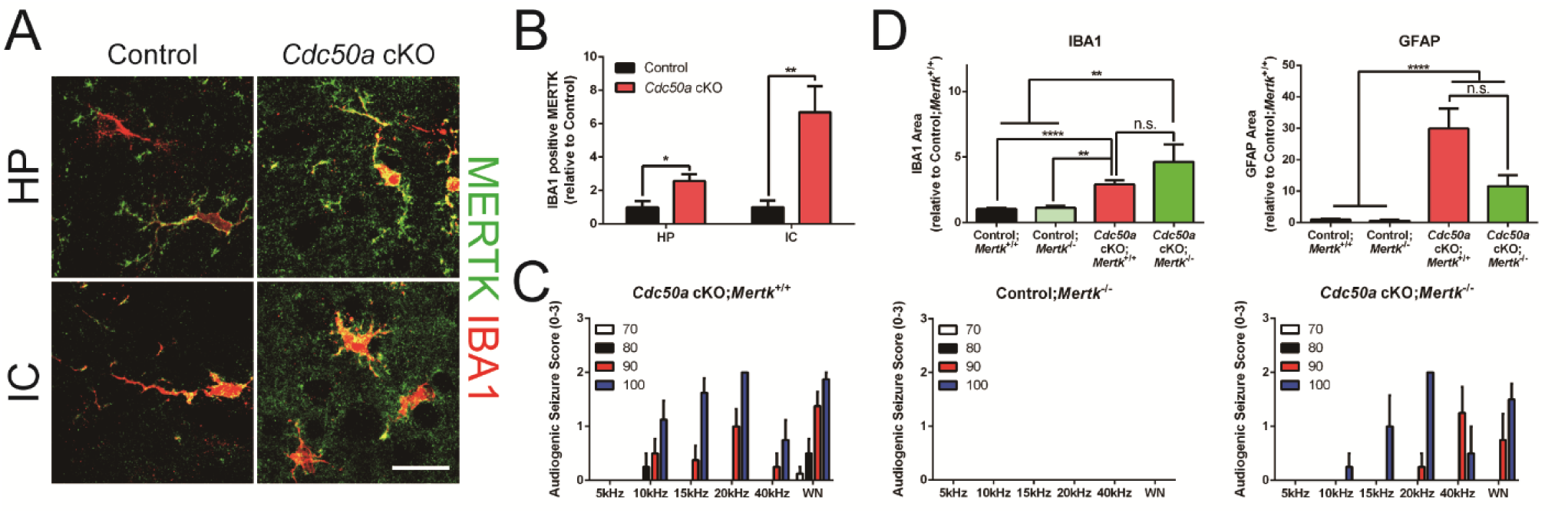
*Mertk* deficiency rescues audiogenic seizure in *Cdc50a* cKO mice. (A) Representative confocal images of MERTK (green) expressing IBA1 (red) positive microglia in control and *Cdc50a* cKO brains. (B) Bar graphs showing the area of MERTK colocalized with IBA1 in control and *Cdc50a* cKO brains. (C) Bar graphs showing audiogenic seizure score in *Cdc50a* cKO;*Mertk*^+/+^, control;*Mertk*^-/-^, and *Cdc50a* cKO;*Mertk*^-/-^ mice. (D) Bar graphs showing area of IBA1 (left) and GFAP (right) in control;*Mertk*^+/+^, *Cdc50a* cKO;*Mertk*^+/+^, control;*Mertk*^-/-^, and *Cdc50a* cKO;*Mertk*^-/-^ brains (IC). *n =* 3 to 6 per group. Bars and error bars indicate mean and SEM, respectively. n.s. = P > 0.05, *P < 0.05, **P < 0.005 or ****P < 0.0001 using *t-*test (B) and Dunnet’s post hoc test (D). Scale bar; 20μm (A).

**Fig. S8:**
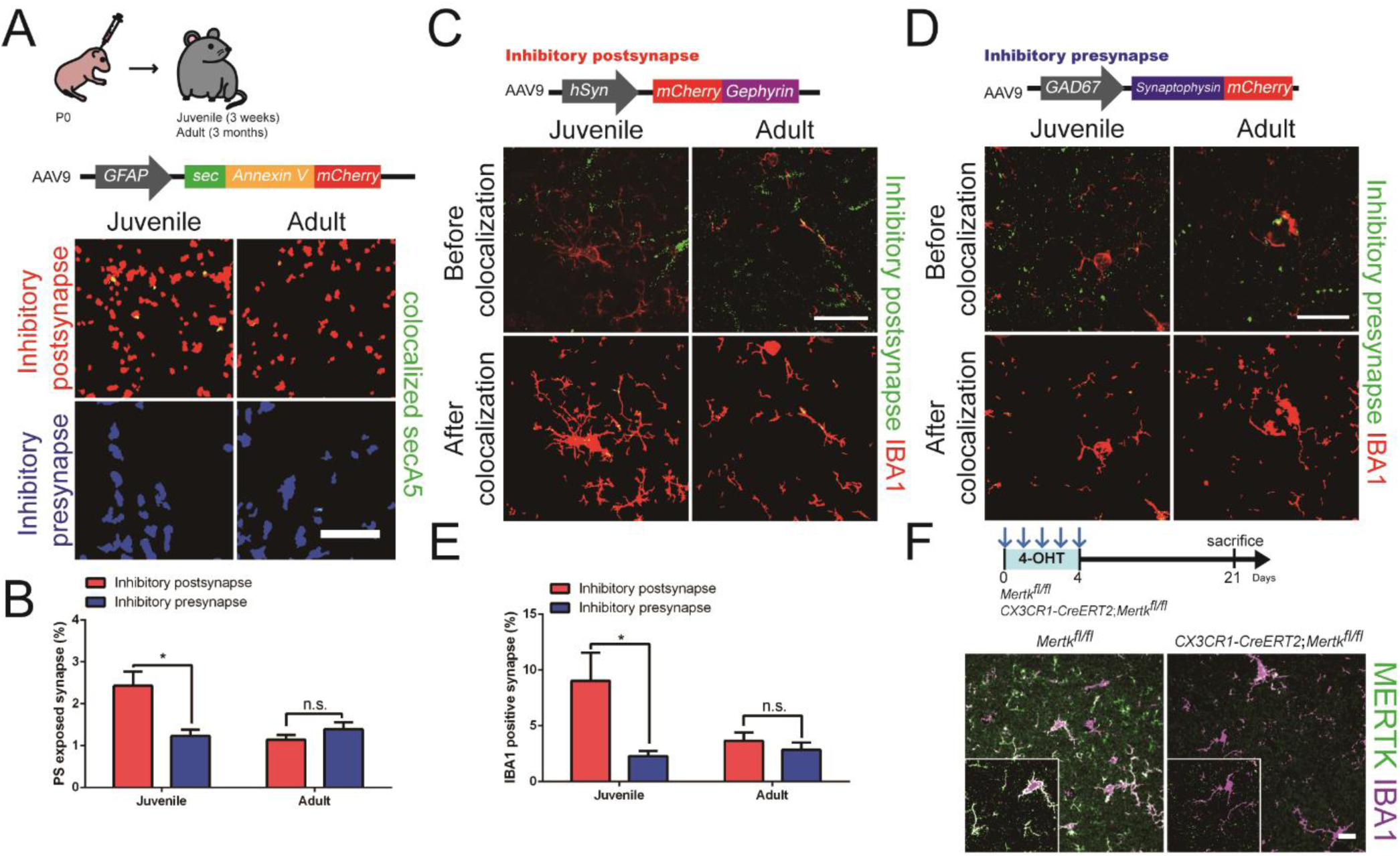
PS-mediated microglial phagocytosis is required for eliminating inhibitory post-synapses in juvenile brains. (A) Representative colocalization images of inhibitory post-synapses (red, upper) and inhibitory pre-synapses (blue, lower) colocalized with secA5 (green) in juvenile and adult WT brains (HP) after injecting AAV9-*GFAP-secA5-mCherry*. (B) Bar graphs showing percentage of inhibitory synapses colocalized with secA5 in juvenile and adult WT brains (HP). (C) Representative confocal (upper) and colocalization (lower) images of inhibitory post-synapses (green) colocalized with IBA1 (red) in juvenile and adult WT brains (HP) after injecting AAV9-*hSyn-mCherry-Gephyrin*. (D) Representative confocal (upper) and colocalization (lower) images of inhibitory pre-synapses (green) colocalized with IBA1 (red) in juvenile and adult WT brains (HP) after injecting AAV9-*GAD67-Synaptophysin-mCherry*. (E) Bar graphs showing percentage of inhibitory synapses colocalized with IBA1 in juvenile and adult WT brains (HP). (F) Representative confocal images of MERTK (green) and IBA1 (magenta) in *Mertk*^*fl/fl*^ and *CX3CR1-CreERT2;Mertk*^*fl/fl*^ mice brains (HP) after injecting 4-OHT. *n =* 4 to 5 per group. Bars and error bars indicate mean and SEM, respectively. n.s. = P > 0.05 or *P < 0.05 using *t-*test (B and E). Scale bar; 10μm (A) or 20μm (C, D, and F).

